## Supplementary material for "Genetic architecture of heritable leaf microbes": Figure S

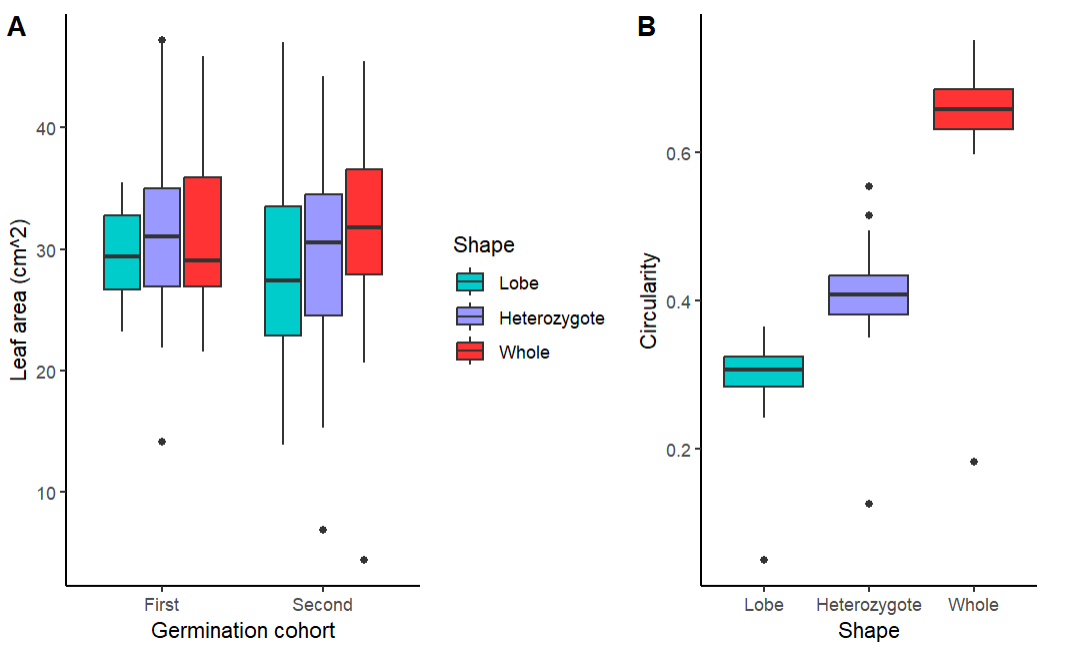

Figure S1. Leaf genotype morphological differences. Leaf area (cm^2^) was not significantly different between collected leaves (A). Circularity significantly differed between *Ipomoea hederacea* leaf shape genotypes (B).

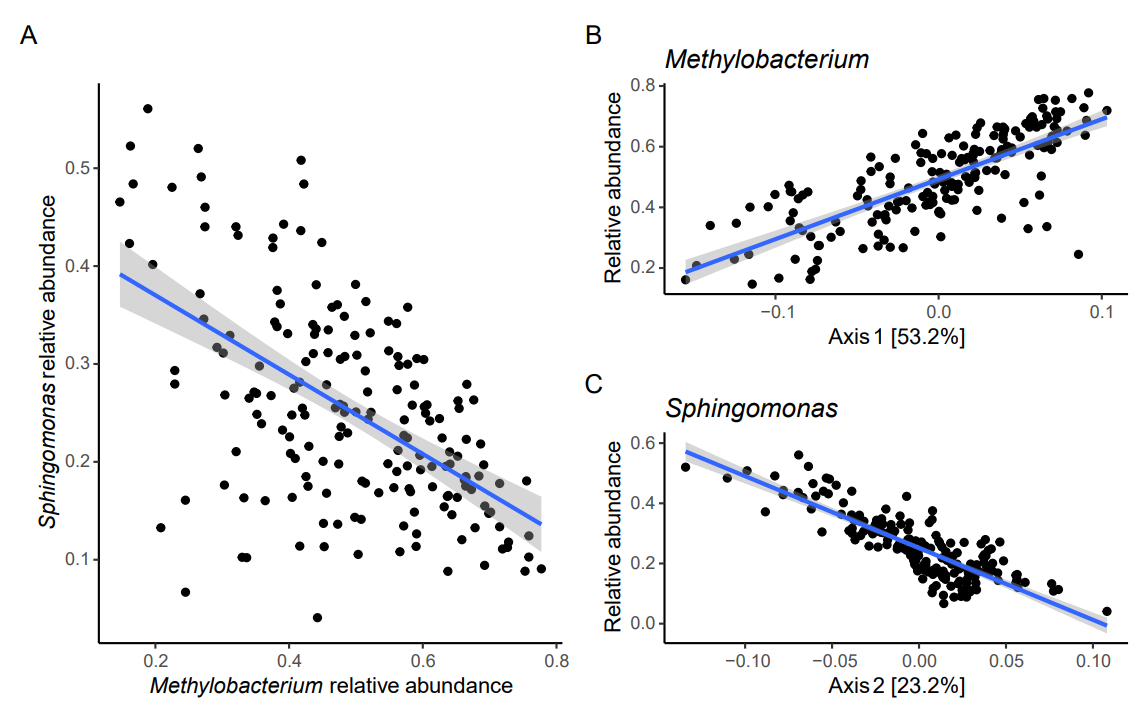

Figure S2. Regressions of relative abundances and PCoA results. The negative correlation in *Methylobacterium* and *Sphingomonas* relative abundances in the phyllosphere (A). The relative abundances of *Methylobacterium* (B) and *Sphingomonas* (C) genera against the first and second axis of the weighted UniFrac PCoA, respectively.

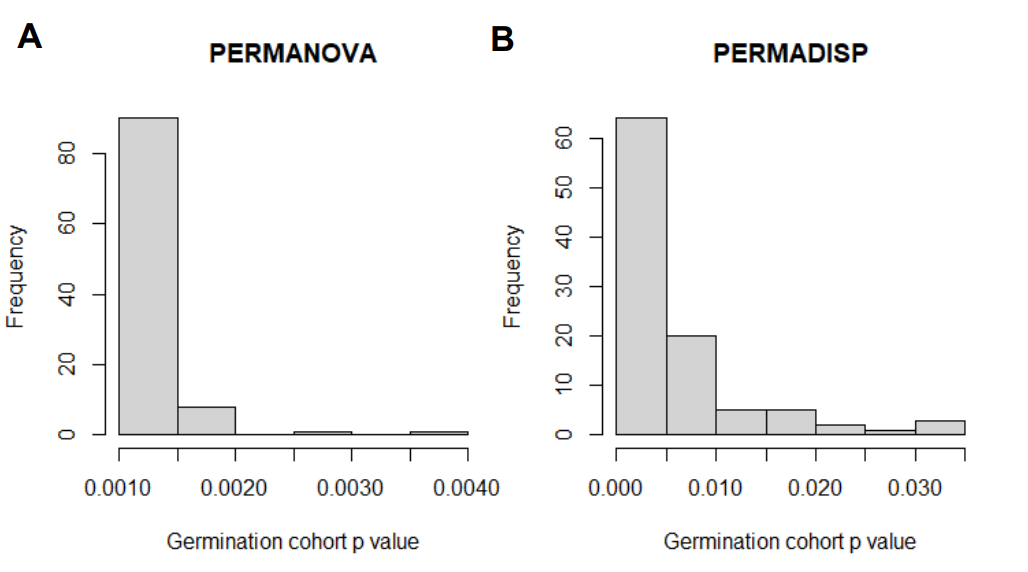

Figure S3. Downsampling to equalize cohort sample sizes (n=77) does not change the significant effect of the germination cohort on microbiome composition (A) and dispersion (B). Histograms show germination cohort p-values from the 100 downsampled weighted UniFrac distance matrices.

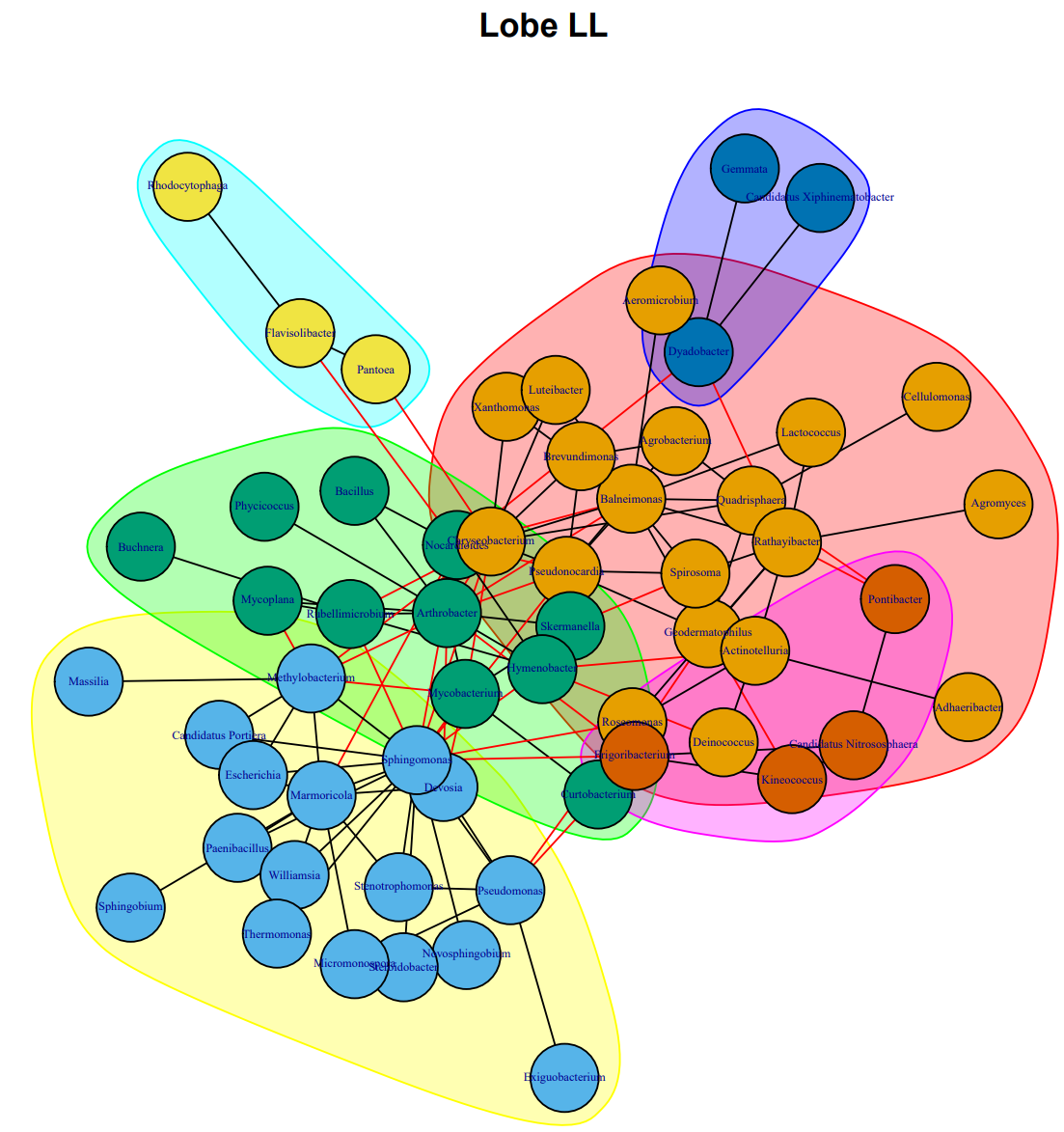

Figure S4. Co-occurrence network of microbial genera in the homozygous lobe leaf shape genotype. Genera are represented by nodes, with node colour and grouping representing a walktrap cluster. Edges indicate a significant correlation between genera, with black edges being a positive correlation and red edges being a negative correlation.

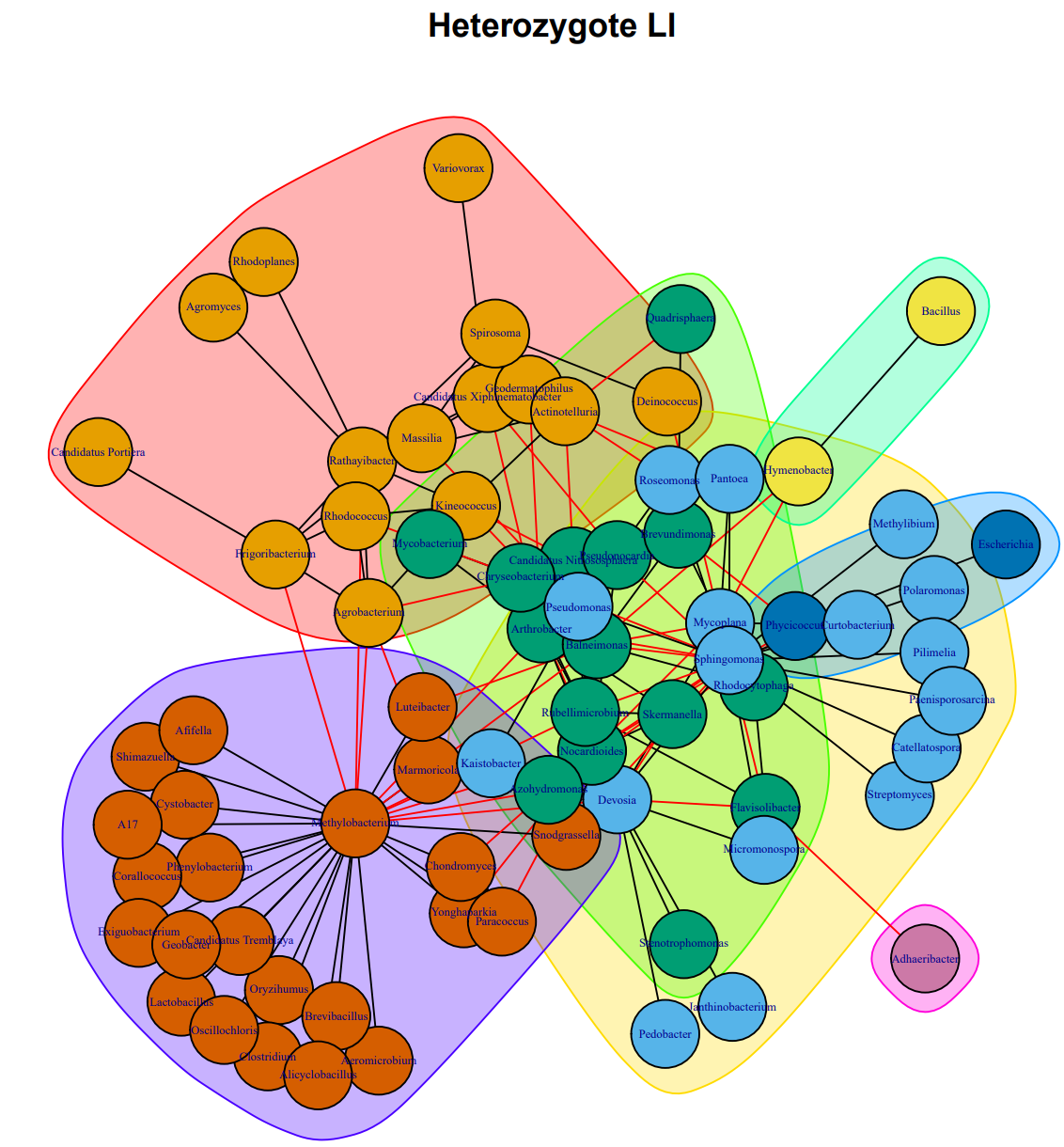

Figure S5. Co-occurrence network of microbial genera in the heterozygous leaf shape genotype. Genera are represented by nodes, with node colour and grouping representing a walktrap cluster. Edges indicate a significant correlation between genera, with black edges being a positive correlation and red edges being a negative correlation.

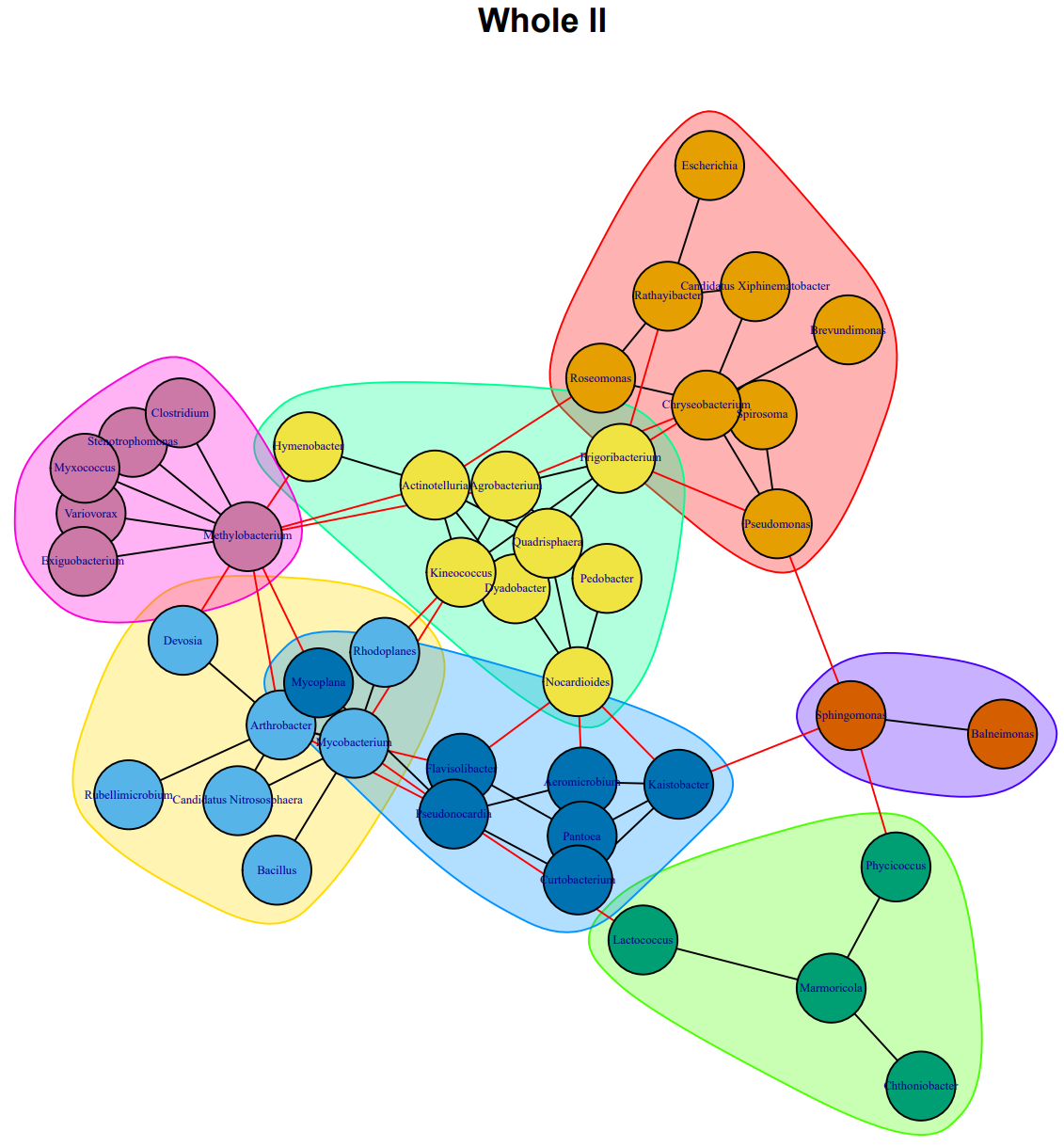

Figure S6. Co-occurrence network of microbial genera in the homozygous whole leaf shape genotype. Genera are represented by nodes, with node colour and grouping representing a walktrap cluster. Edges indicate a significant correlation between genera, with black edges being a positive correlation and red edges being a negative correlation.

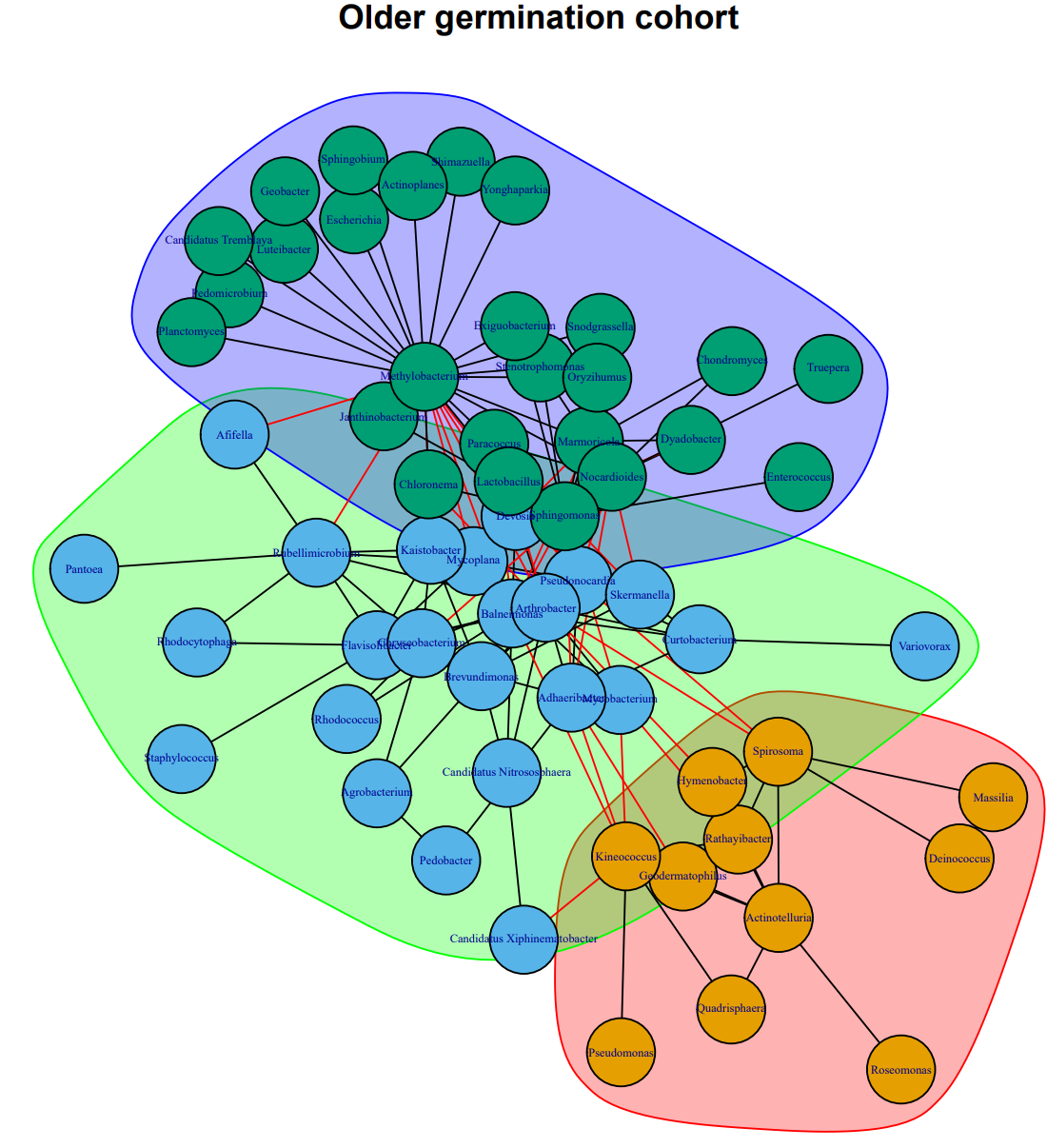

Figure S7. Co-occurrence network of microbial genera in the older germination cohort. Genera are represented by nodes, with node colour and grouping representing a walktrap cluster. Edges indicate a significant correlation between genera, with black edges being a positive correlation and red edges being a negative correlation.

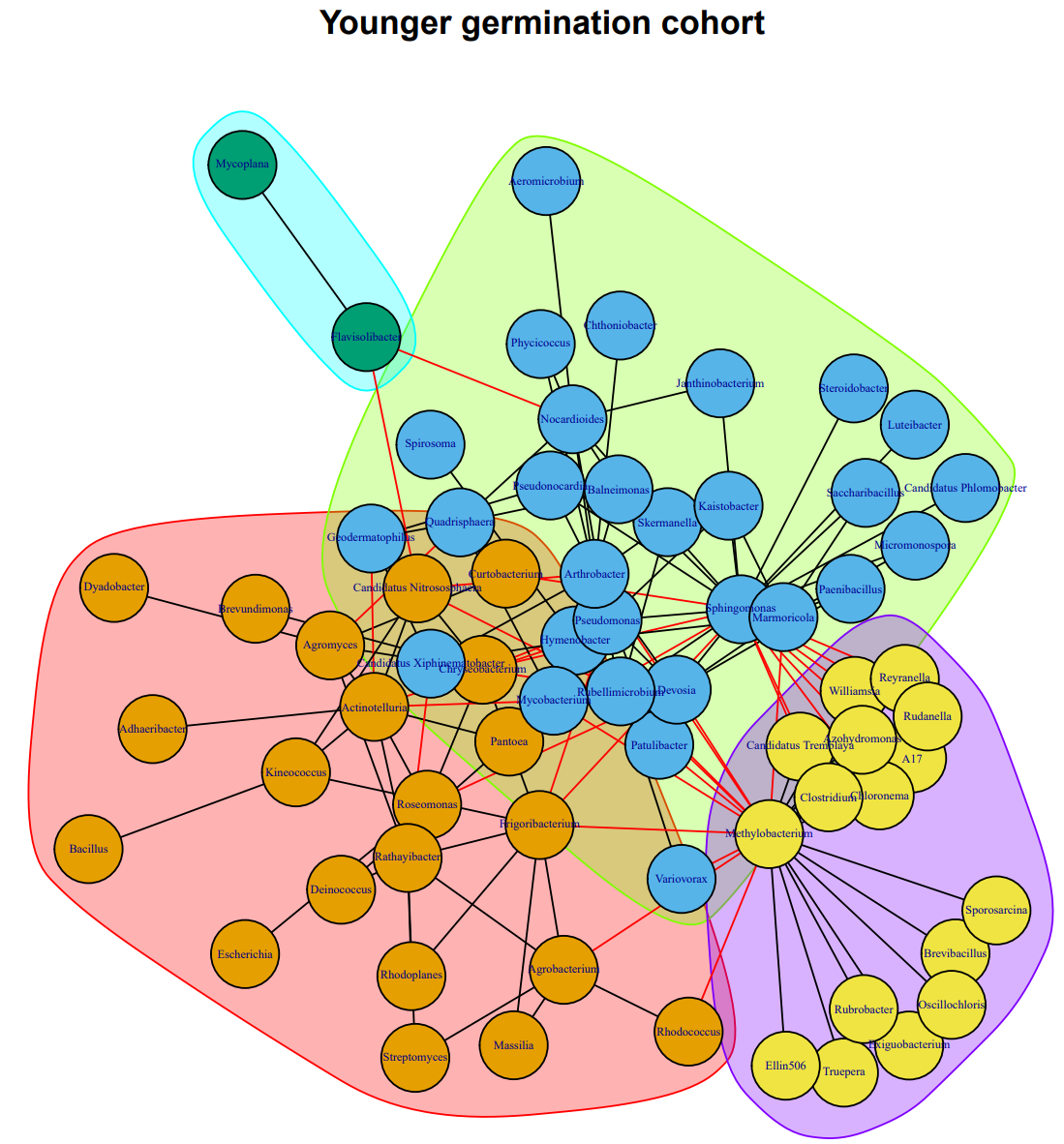

Figure S8. Co-occurrence network of microbial genera in the younger germination cohort. Genera are represented by nodes, with node colour and grouping representing a walktrap cluster. Edges indicate a significant correlation between genera, with black edges being a positive correlation and red edges being a negative correlation.

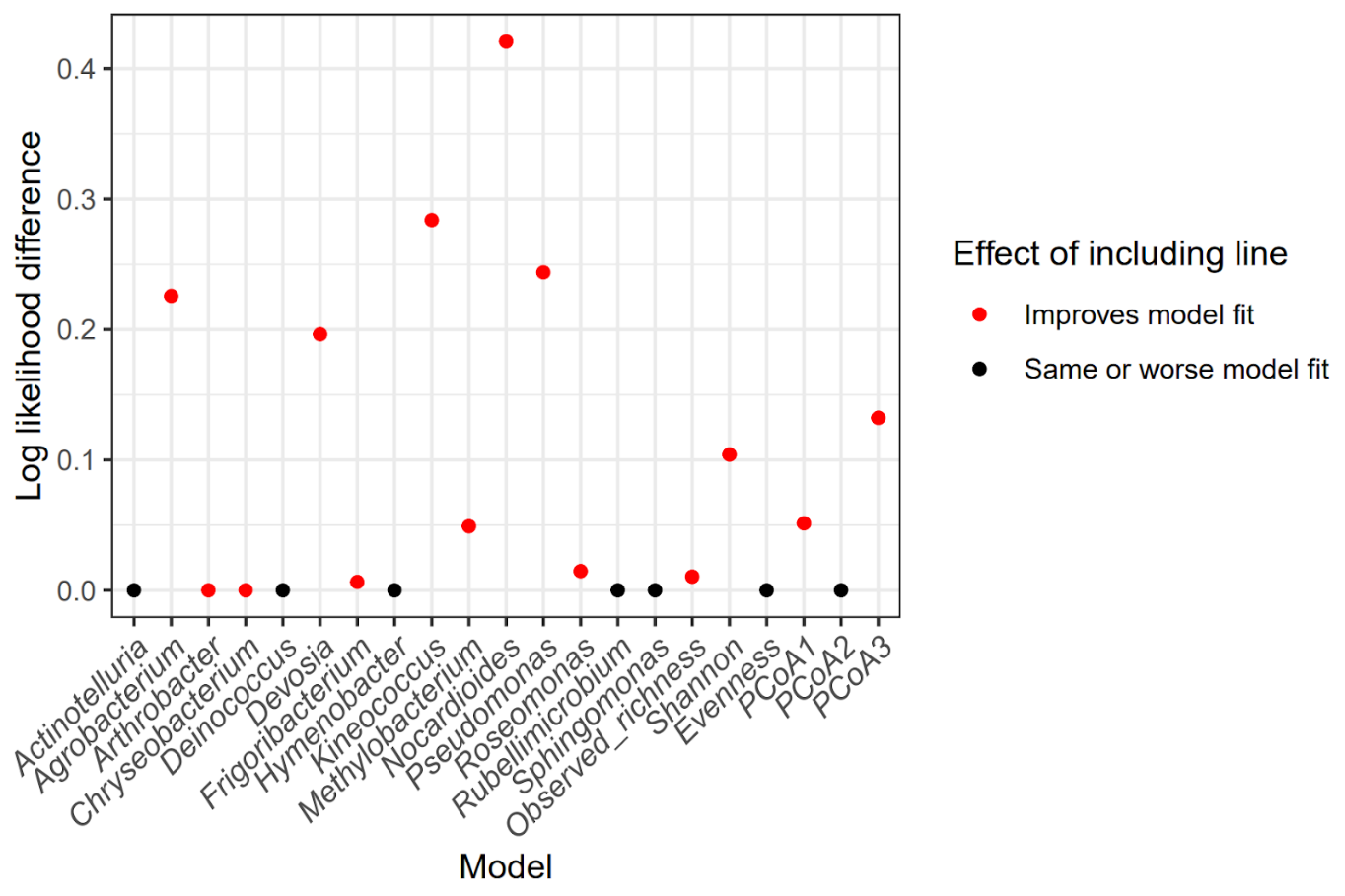

Figure S9. Log-likelihood difference between broad-sense heritability models including line as a random effect or not. Red color indicates a significant improvement by including line, while black indicates the same or slightly worse fit by including line.
