## Supplementary material for "Genetic architecture of heritable leaf microbes": Table S

Table S1. Pearson’s chi-squared test for differences in leaf shape individual counts between germination cohorts.

|  | **Leaf shape individuals (no.)** | | |
| --- | --- | --- | --- |
| *Predictors* | *X^2^ value* | *df* | *p* |
| Germination cohort | 0.311 | 2 | 0.856 |

Table S2. Linear mixed effect model results for leaf morphology. Wald *X*^2^ values and p values are from type III ANOVAs.

|  | **Leaf surface area (cm^2^)** | | | **Leaf circularity** | | |
| --- | --- | --- | --- | --- | --- | --- |
| *Predictors* | *Wald X^2^ value* | *df* | *p* | *Wald X^2^ value* | *df* | *p* |
| Intercept | 203 | 1 | **<0.001** | 152e1 | 1 | **<0.001** |
| Leaf shape | 4.15 | 2 | 0.125 | 102e1 | 2 | **<0.001** |
| Germination cohort | 2.22 | 1 | 0.137 | 6.9e-3 | 1 | 0.934 |

Table S3. PERMANOVA and PERMADISP results for microbiome composition, using rarefied weighted Unifrac distances.

|  | **PERMANOVA: Weighted Unifrac** | | | **PERMADISP: Weighted Unifrac** | | |
| --- | --- | --- | --- | --- | --- | --- |
| *Predictors* | *F* | *df* | *p* | *F* | *df* | *p* |
| Leaf shape | 1.83 | 2, 182 | 0.079 | - | - | - |
| Germination cohort | 10.4 | 1, 182 | **0.001** | 10.4 | 1, 182 | **0.001** |

Table S4. Linear mixed effect model results for heritability of community phenotypes, including leaf genotype and germination cohort as fixed effects and line and block as random effects. Wald *X*^2^ values, F values, and p values are from type III ANOVAs. Random effect results are from the linear model directly.

|  | **log10(Observed richness)** | | | **Shannon diversity** | | | **PCoA1** | | | **PCoA2** | | | **PCoA3** | | | **Evenness** | | |
| --- | --- | --- | --- | --- | --- | --- | --- | --- | --- | --- | --- | --- | --- | --- | --- | --- | --- | --- |
| *Fixed Effects* | *Wald X^2^ value* | *df* | *p* | *Wald X^2^ value* | *df* | *p* | *Wald X^2^ value* | *df* | *p* | *Wald X^2^ value* | *df* | *p* | *Wald X^2^ value* | *df* | *p* | *Wald X^2^ value* | *df* | *p* |
| Intercept | 285e1 | 1 | **<0.001** | 143e1 | 1 | **<0.001** | 1.03 | 1 | 0.311 | 2.02 | 1 | 0.156 | 5.85 | 1 | **0.016** | 533e1 | 1 | <0.001 |
| Leaf shape | 3.77 | 2 | 0.152 | 3.97 | 2 | 0.137 | 3.32 | 2 | 0.190 | 0.362 | 2 | 0.835 | 4.85 | 2 | 0.089 | 1.42 | 2 | 0.493 |
| Germination cohort | 10.1 | 1 | **0.001** | 3.33 | 1 | 0.068 | 18.9 | 1 | **<0.001** | 6.71 | 1 | **0.010** | 2.98 | 1 | 0.084 | 2.96 | 1 | 0.085 |
| *Random Effects* | *Variance* | *Std. Dev.* | | *Variance* | *Std. Dev.* | | *Variance* | *Std. Dev.* | | *Variance* | *Std. Dev.* | | *Variance* | *Std. Dev.* | | *Variance* | *Std. Dev.* | |
| Line | 2.28e-3 | 4.78e-2 | | 6.89e-3 | 0.083 | | 7.61e-5 | 8.73e-3 | | 0.0 | 0.0 | | 2.30e-5 | 4.80e-3 | | 0.0 | 0.0 | |
| Block | 2.45e-4 | 1.56e-2 | | 6.10e-3 | 0.078 | | 2.53e-4 | 0.016 | | 0.0 | 0.0 | | 6.43e-7 | 8.02e-4 | | 1.29e-4 | 0.011 | |
| Residual | 0.195 | 0.442 | | 0.184 | 0.429 | | 2.62e-3 | 0.051 | | 1.35e-3 | 0.037 | | 4.56e-4 | 0.021 | | 2.60e-3 | 0.051 | |

Table S5. SparCC co-occurrence network properties, using only false discovery rate corrected significant correlations.

| *Network property* | **Network** | | | | |
| --- | --- | --- | --- | --- | --- |
|  | *Lobe LL* | *Hetero Ll* | *Heart ll* | *Germ. cohort 1* | *Germ. cohort 2* |
| Node number | 57 | 75 | 43 | 61 | 68 |
| Edge number | 122 | 151 | 68 | 149 | 140 |
| Walktrap cluster number | 6 | 6 | 7 | 3 | 4 |
| Global clustering coefficient | 0.226 | 0.163 | 0.142 | 0.240 | 0.118 |
| Modularity | 0.413 | 0.421 | 0.477 | 0.349 | 0.354 |
| Top 5 ‘hub’ genera | *1.Sphingomonas*  *2.Chryseobacterium*  *3.Arthrobacter*  *4.Nocardioides*  *5.Balneimonas* | *1.Methylobacterium*  *2.Sphingomonas*  *3.Nocardioides*  *4.Arthrobacter*  *5.Devosia* | *1.Methylobacterium*  *2.Arthrobacter*  *3.Kineococcus*  *4.Actinotelluria*  *5.Agrobacterium* | *1.Arthrobacter*  *2.Methylobacterium*  *3.Balneimonas*  *4.Pseudonocardia*  *5.Nocardioides* | *1.Sphingomonas*  *2.Methylobacterium*  *3.Arthrobacter*  *4.Rubellimicrobium*  *5.Devosia* |
| Number of positive correlations | 62 | 93 | 43 | 88 | 83 |
| Number of negative correlations | 60 | 58 | 25 | 61 | 57 |

Table S6. Linear mixed effect model results for comparing the magnitude of effect of matriline and leaf shape on community phenotypes. The models included germination cohort as a fixed effect, and line, leaf genotype, and block as random effects. Since the models’ purpose is solely to compare the magnitude of effect, only variance explained by random effects is shown.

|  | **log10(Observed richness)** | | **Shannon diversity** | | **PCoA1** | | **PCoA2** | | **PCoA3** | |
| --- | --- | --- | --- | --- | --- | --- | --- | --- | --- | --- |
| *Random Effects* | *Variance* | *Std. Dev.* | *Variance* | *Std. Dev.* | *Variance* | *Std. Dev.* | *Variance* | *Std. Dev.* | *Variance* | *Std. Dev.* |
| Line | 1.09e-3 | 0.033 | 6.53e-3 | 0.080 | 9.74e-5 | 9.87e-3 | 0.00 | 0.00 | 1.63e-5 | 4.04e-3 |
| Block | 8.55e-4 | 0.029 | 6.60e-3 | 0.081 | 2.84e-4 | 0.017 | 2.38e-6 | 1.54e-3 | 1.40e-12 | 1.18e-6 |
| Leaf shape | 2.96e-3 | 0.054 | 3.37e-3 | 0.058 | 3.31e-5 | 5.75e-3 | 0.00 | 0.00 | 1.13e-5 | 3.37e-3 |
| Residual | 0.196 | 0.442 | 0.184 | 0.429 | 2.59e3 | 0.051 | 1.33e-3 | 0.037 | 4.62e-4 | 2.15e-2 |

Table S7. Linear mixed effect model results for heritability of bacterial genera, including leaf genotype and germination cohort as fixed effects and line and block as random effects. Wald *X*^2^ values, F values, and p values are from type III ANOVAs. Random effect results are from the linear model directly.

|  | ***Actinotelluria*** | | | ***Agrobacterium*** | | | ***Arthrobacter*** | | | ***Chryseobacterium*** | | | ***Deinococcus*** | | |
| --- | --- | --- | --- | --- | --- | --- | --- | --- | --- | --- | --- | --- | --- | --- | --- |
| *Fixed Effects* | *Wald X^2^ value* | *df* | *p* | *Wald X^2^ value* | *df* | *p* | *Wald X^2^ value* | *df* | *p* | *Wald X^2^ value* | *df* | *p* | *Wald X^2^ value* | *df* | *p* |
| Intercept | 30.1 | 1 | **<0.001** | 4.81 | 1 | **0.028** | 1.95 | 1 | 0.162 | 14.9 | 1 | **<0.001** | 7.45 | 1 | **0.006** |
| Leaf shape | 2.18 | 2 | 0.337 | 11.4 | 2 | **0.003** | 0.161 | 2 | 0.923 | 0.817 | 2 | 0.665 | 1.58 | 2 | 0.454 |
| Germination cohort | 6.68 | 1 | **0.010** | 0.106 | 1 | 0.744 | 7.67 | 1 | **0.006** | 7.63 | 1 | **0.006** | 12.0 | 1 | **<0.001** |
| *Random Effects* | *Variance* | *Std. Dev.* | | *Variance* | *Std. Dev.* | | *Variance* | *Std. Dev.* | | *Variance* | *Std. Dev.* | | *Variance* | *Std. Dev.* | |
| Line | 0.00 | 0.00 | | 0.100 | 0.317 | | 0.00 | 0.00 | | 0.00 | 0.00 | | 0.00 | 0.00 | |
| Block | 0.012 | 0.108 | | 0.035 | 0.186 | | 0.290 | 0.539 | | 0.287 | 0.536 | | 0.00 | 0.00 | |
| Residual | 2.01 | 1.42 | | 1.89 | 1.37 | | 2.40 | 1.55 | | 3.19 | 1.77 | | 2.49 | 1.58 | |

|  | ***Devosia*** | | | ***Frigoribacterium*** | | | ***Hymenobacter*** | | | ***Kineococcus*** | | | ***Methylobacterium*** | | |
| --- | --- | --- | --- | --- | --- | --- | --- | --- | --- | --- | --- | --- | --- | --- | --- |
| *Fixed Effects* | *Wald X^2^ value* | *df* | *p* | *Wald X^2^ value* | *df* | *p* | *Wald X^2^ value* | *df* | *p* | *Wald X^2^ value* | *df* | *p* | *Wald X^2^ value* | *df* | *p* |
| Intercept | 31.8 | 1 | **<0.001** | 5.44e-3 | 1 | 0.941 | 100 | 1 | **<0.001** | 30.1 | 1 | **<0.001** | 583 | 1 | <0.001 |
| Leaf shape | 1.69 | 2 | 0.430 | 1.31 | 2 | 0.518 | 4.36 | 2 | 0.113 | 2.86 | 2 | 0.239 | 7.03 | 2 | **0.030** |
| Germination cohort | 0.802 | 1 | 0.371 | 6.06 | 1 | **0.014** | 2.27 | 1 | 0.132 | 0.031 | 1 | 0.860 | 3.83 | 1 | **0.050** |
| *Random Effects* | *Variance* | *Std. Dev.* | | *Variance* | *Std. Dev.* | | *Variance* | *Std. Dev.* | | *Variance* | *Std. Dev.* | | *Variance* | *Std. Dev.* | |
| Line | 0.084 | 0.290 | | 0.019 | 0.138 | | 0.00 | 0.00 | | 0.131 | 0.362 | | 0.021 | 0.146 | |
| Block | 0.011 | 0.104 | | 0.064 | 0.253 | | 0.00 | 0.00 | | 0.038 | 0.195 | | 9.07e-3 | 0.095 | |
| Residual | 1.73 | 1.32 | | 2.34 | 1.53 | | 1.55 | 1.24 | | 2.10 | 1.45 | | 0.910 | 0.954 | |

|  | ***Nocardioides*** | | | ***Pseudomonas*** | | | ***Roseomonas*** | | | ***Rubellimicrobium*** | | | ***Sphingomonas*** | | |
| --- | --- | --- | --- | --- | --- | --- | --- | --- | --- | --- | --- | --- | --- | --- | --- |
| *Fixed Effects* | *Wald X^2^ value* | *df* | *p* | *Wald X^2^ value* | *df* | *p* | *Wald X^2^ value* | *df* | *p* | *Wald X^2^ value* | *df* | *p* | *Wald X^2^ value* | *df* | *p* |
| Intercept | 0.331 | 1 | 0.565 | 2.51e-2 | 1 | 0.874 | 11.1 | 1 | **<0.001** | 34.0 | 1 | **<0.001** | 541 | 1 | **<0.001** |
| Leaf shape | 7.61 | 2 | **0.022** | 1.58 | 2 | 0.455 | 1.19 | 2 | 0.553 | 3.20 | 2 | 0.202 | 7.78 | 2 | **0.020** |
| Germination cohort | 0.032 | 1 | 0.858 | 16.6 | 1 | **<0.001** | 11.5 | 1 | **<0.001** | 3.23 | 1 | 0.072 | 4.79 | 1 | **0.029** |
| *Random Effects* | *Variance* | *Std. Dev.* | | *Variance* | *Std. Dev.* | | *Variance* | *Std. Dev.* | | *Variance* | *Std. Dev.* | | *Variance* | *Std. Dev.* | |
| Line | 0.212 | 0.461 | | 0.118 | 0.343 | | 0.021 | 0.144 | | 0.00 | 0.00 | | 0.00 | 0.00 | |
| Block | 0.030 | 0.173 | | 0.00 | 0.00 | | 0.015 | 0.122 | | 0.204 | 0.452 | | 0.00 | 0.00 | |
| Residual | 2.64 | 1.62 | | 2.18 | 1.48 | | 1.56 | 1.25 | | 2.39 | 1.55 | | 0.706 | 0.840 | |

Table S8. Linear mixed effect model results for comparing the magnitude of effect of matriline and leaf shape on genera. The models included germination cohort as a fixed effect, and line, leaf genotype, and block as random effects. Since the models’ purpose is solely to compare the magnitude of effect, only variance explained by random effects is shown.

|  | ***Actinotelluria*** | | ***Agrobacterium*** | | ***Arthrobacter*** | | ***Chryseobacterium*** | | ***Deinococcus*** | |
| --- | --- | --- | --- | --- | --- | --- | --- | --- | --- | --- |
| *Random Effects* | *Variance* | *Std. Dev.* | *Variance* | *Std. Dev.* | *Variance* | *Std. Dev.* | *Variance* | *Std. Dev.* | *Variance* | *Std. Dev.* |
| Line | 4.64e-3 | 0.068 | 0.114 | 0.338 | 0.00 | 0.00 | 0.00 | 0.00 | 0.00 | 0.00 |
| Block | 0.018 | 0.133 | 0.034 | 0.185 | 0.284 | 0.533 | 0.299 | 0.547 | 3.31e-3 | 0.057 |
| Leaf shape | 0.00 | 0.00 | 0.122 | 0.350 | 0.00 | 0.00 | 0.00 | 0.00 | 0.00 | 0.00 |
| Residual | 2.00 | 1.41 | 1.87 | 1.37 | 2.38 | 1.54 | 3.17 | 1.78 | 2.48 | 1.57 |

|  | ***Devosia*** | | ***Frigoribacterium*** | | ***Hymenobacter*** | | ***Kineococcus*** | | ***Methylobacterium*** | |
| --- | --- | --- | --- | --- | --- | --- | --- | --- | --- | --- |
| *Random Effects* | *Variance* | *Std. Dev.* | *Variance* | *Std. Dev.* | *Variance* | *Std. Dev.* | *Variance* | *Std. Dev.* | *Variance* | *Std. Dev.* |
| Line | 0.064 | 0.253 | 3.97e-3 | 0.063 | 0.00 | 0.00 | 0.108 | 0.329 | 0.021 | 0.145 |
| Block | 0.015 | 0.124 | 0.063 | 0.250 | 5.54e-10 | 2.35e-5 | 0.043 | 0.208 | 0.010 | 0.101 |
| Leaf shape | 0.00 | 0.00 | 0.00 | 0.00 | 2.75e-2 | 0.166 | 0.014 | 0.116 | 0.033 | 0.181 |
| Residual | 1.75 | 1.32 | 2.35 | 1.53 | 1.55 | 1.24 | 2.12 | 1.46 | 0.909 | 0.954 |

|  | ***Nocardioides*** | | ***Pseudomonas*** | | ***Roseomonas*** | | ***Rubellimicrobium*** | | ***Sphingomonas*** | |
| --- | --- | --- | --- | --- | --- | --- | --- | --- | --- | --- |
| *Random Effects* | *Variance* | *Std. Dev.* | *Variance* | *Std. Dev.* | *Variance* | *Std. Dev.* | *Variance* | *Std. Dev.* | *Variance* | *Std. Dev.* |
| Line | 0.195 | 0.441 | 9.70e-2 | 0.311 | 8.33e-3 | 0.091 | 0.00 | 0.00 | 0.00 | 0.00 |
| Block | 0.035 | 0.188 | 2.01e-9 | 4.48e-5 | 0.017 | 0.129 | 0.209 | 0.457 | 0.00 | 0.00 |
| Leaf shape | 0.110 | 0.331 | 1.50e-8 | 1.22e-4 | 0.00 | 0.00 | 0.019 | 0.136 | 0.027 | 0.166 |
| Residual | 2.65 | 1.63 | 2.19 | 1.48 | 1.57 | 1.25 | 2.39 | 1.55 | 0.705 | 0.840 |

Table S9. Heritability patterns of microbes across studies. For comparison to morning glory, the selected studies focus on plant microbes but also include estimates in animals.

| **Study** | **Host and compartment** | **Metric** | **Range of heritability** | **Mean heritability** |
| --- | --- | --- | --- | --- |
| Walters et al. 2018 | Maize rhizosphere | Core OTU abundance | H^2^=0.15-0.25 | NA |
| Peiffer et al. 2013 | Maize rhizosphere | OTU richness | H^2^=0.18-0.19 | H^2^=0.19 |
| Peiffer et al. 2013 | Maize rhizosphere | Weighted Unifrac 𝛽-diversity | H^2^=0.07-0.15 | H^2^=0.08 |
| Peiffer et al. 2013 | Maize rhizosphere | Unweighted Unifrac 𝛽-diversity | H^2^=0.05-0.06 | H^2^=0.05 |
| Wallace et al. 2018 | Maize phyllosphere | OTU and higher taxonomic groups abundance | h^2^=0-0.41 | NA |
| Wallace et al. 2018 | Maize phyllosphere | ⍺-diversity | h^2^=0-0.05 | h^2^≅0 |
| Horton et al. 2014 | *Arabidopsis thaliana* phyllosphere | PCA Community eigenvectors | H^2^=0.01-0.16 | H^2^=0.06 |
|  |  | OTU richness | NA | H^2^≅0.46 |
| Wagner et al. 2016 | *Boechera stricta* rhizosphere and phyllosphere | OTU abundance | H^2^=0-1.0 | H^2^≤0.10 |
| Deng et al. 2021 | Sorghum rhizosphere | OTU abundance | H^2^=0-0.66 | H^2^=0.10 |
| Edwards et al. 2023 | Switchgrass rhizosphere | ASV abundance | V_A_≅0-0.40 | V_A_ ≅0.10 |
| Edwards et al. 2023 | Switchgrass rhizosphere | Genus abundances | V_A_≅0-0.20 | V_A_ ≅0.05 |
| Schweitzer et al. 2008 | *Populus* tree rhizospheres | Bray-Curtis dissimilarity | H^2^=0.30-1.0 | H^2^=0.70 |
| Wallace et al. 2019 | Dairy cow rumen | Core OTU abundance | h^2^=0.20-0.60 | NA |
| Grieneisen et al. 2021 | Baboon gut | ASV abundance and community phenotypes | h^2^=0-0.21 | h^2^=0.07 |
| Goodrich et al. 2014 | Human gut | OTU abundance | H^2^≅0-0.35 | NA |

Table S10. Heritability of *Arabidopsis thaliana* traits using different crossing designs.

| **Study** | **Plant material** | **Phenotype** | **Environment** | **Heritability** |
| --- | --- | --- | --- | --- |
| Brachi et al. 2010 | Natural accessions (n=197) | Flowering time | Field | H^2^=0.97 |
|  | RIL parents (n=14) | Flowering time | Field | H^2^=0.92 |
|  | RIL families (n=13) | Flowering time | Field | H^2^=0.71-0.94, mean 0.81 |
| Weinig et al. 2002 | RILs (n=98) | Date of bolting | Field | H^2^=0.10-0.22 |
|  |  |  | Growth chamber | H^2^=0.44-0.49 |
| Méndez-Vigo et al. 2012 | Natural accessions (n=8) | Flowering time | Field | H^2^= 0-0.77, mean 0.36 |
|  |  |  | Glasshouse | H^2^=0.08-0.71, mean 0.42 |
|  |  | Number of rosettes | Field | H^2^=0.23-0.69, mean 0.48 |
|  |  | Leaf number | Glasshouse | H^2^=0.20-0.84, mean 0.58 |
|  |  | Number of fruiting plants | Field | H^2^=0.0-0.46, mean 0.22 |
| Salomé et al. 2011 | Natural accessions (n=18) | Flowering time | Growth chamber | Approximate H^2^=0.15-0.95, Approximate mean of 0.55 |
|  | F_2_ populations from the accessions (n=17) | Flowering time | Growth chamber | H^2^=0.10-0.64, mean 0.39 |
| Sasaki et al. 2015 | Natural inbred lines (n=173) | Flowering time | Greenhouse 10℃ | H^2^=0.97 |
|  |  |  | Greenhouse 16℃ | H^2^=0.91 |
| Lempe et al. 2005 | Natural accessions (n=150) | Flowering time | 16 °C long days | H^2^=0.81 |
|  |  |  | 23 °C long days | H^2^=0.85 |
|  |  |  | 16 °C long days after 5-week vernalization | H^2^=0.70 |
|  |  |  | 23 °C short days | H^2^=0.71 |
|  |  | Leaf number | 16 °C long days | H^2^=0.90 |
|  |  |  | 23 °C long days | H^2^=0.89 |
|  |  |  | 16 °C long days after 5-week vernalization | H^2^=0.84 |
|  |  |  | 23 °C short days | H^2^=0.87 |
